# Electrobiocorrosion by Microbes without Outer-Surface Cytochromes

**DOI:** 10.1101/2023.07.26.550717

**Authors:** Dawn E. Holmes, Trevor L. Woodard, Jessica A. Smith, Florin Musat, Derek R. Lovley

**Author notes:** Correspondence: Dawn Holmes, Running title: Electrobiocorrosion by methanogens and acetogens.

## Abstract

Anaerobic microbial corrosion of iron-containing metals causes extensive economic damage. Some microbes are capable of direct metal-to-microbe electron transfer (electrobiocorrosion), but the prevalence of electrobiocorrosion among diverse methanogens and acetogens is poorly understood because of a lack of tools for their genetic manipulation. Previous studies have suggested that respiration with 316L stainless steel as the electron donor is indicative of electrobiocorrosion because, unlike pure Fe^0^, 316L stainless steel does not abiotically generate H_2_ as an intermediary electron carrier. Here we report that all of the methanogens (*Methanosarcina vacuolata*, *Methanothrix soehngenii*, and *Methanobacterium* strain IM1) and acetogens (*Sporomusa ovata*, *Clostridium ljungdahlii*) evaluated respired with pure Fe^0^ as the electron donor, but only *M. vacuolata*, *Mx soehngenii*, and *S. ovata* were capable of stainless steel electrobiocorrosion. The electrobiocorrosive methanogens required acetate as an additional energy source in order to produce methane from stainless steel. Co-cultures of *S. ovata* and *Mx. soehngenii* demonstrated how acetogens can provide acetate to methanogens during corrosion. Not only was *Methanobacterium* strain IM1 not capable of electrobiocorrosion, but it also did not accept electrons from *Geobacter metallireducens*, an effective electron-donating partner for direct interspecies electron transfer to all methanogens that can directly accept electrons from Fe^0^. The finding that *M. vacuolata*, *Mx. soehngenii*, and *S. ovata* are capable of electrobiocorrosion, despite a lack of the outer-surface *c*-type cytochromes previously found to be important in other electrobiocorrosive microbes, demonstrates that there are multiple microbial strategies for making electrical contact with Fe^0^.

**Impact Statement:** Understanding how anaerobic microbes receive electrons from Fe^0^ is likely to lead to novel strategies for mitigating the corrosion of iron-containing metals, which has an enormous economic impact. Electrobiocorrosion, is a relatively recently recognized corrosion mechanism. It was previously demonstrated in pure cultures when Fe^0^ oxidation was inhibited by deletion of genes for outer-surface *c*-type cytochromes known to be involved in other forms of extracellular electron exchange. However, many methanogens and acetogens lack obvious outer-surface electrical connections and are difficult to genetically manipulate. The study reported here provides an alternative approach to evaluating whether microbes are capable of electrobiocorrosion that does not require genetic manipulation. The results indicate that *Methanobacterium* strain IM1, is not electrobiocorrosive, in contrast to previous speculation. However, some methanogens and acetogens without known outer-surface *c*-type cytochromes do appear to be capable of electrobiocorrosion, suggesting that this corrosion mechanism may be more widespread than previously thought.

## Introduction

The costs of microbial corrosion dwarf the negative economic impacts of all other problematic biofilm damage combined, including biomedical and environmental (1). Corrosion of iron-based materials has a particularly significant impact on a variety of industries including tanks and pipes used for gas production and storage, wastewater treatment plants, electric power generation facilities, water distribution networks, and nuclear waste storage facilities (2–4). Understanding the mechanisms of microbial metal corrosion is expected to aid in the development of strategies for detecting microbial corrosion and its prevention.

Metallic iron corrodes when Fe^0^ is oxidized to Fe^+2^:

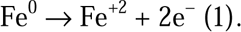

Anaerobic processes are responsible for most microbial corrosion (2–4). The most intensively studied anaerobic corrosion processes are those that involve the electrons released from Fe^0^ reducing protons to generate H_2_. The most basic reaction is:

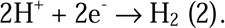

Microbial production of carbon dioxide or organic acids, generate localized low pH zones that favor H_2_ production (5–7), as do extracellular hydrogenases that are either released from lysed cells (8–10), or specifically designed for extracellular release (11). Furthermore, sulfide generated from microbial sulfate reduction reacts with Fe^0^ to produce H_2_ (12):

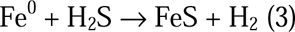

Iron sulfide deposits can also stimulate proton reduction (reaction 2) (2). Microbial consumption of the H_2_ is expected to make reactions 2 and 3 more thermodynamically favorable and accelerate their reaction rates (13).

A more recently recognized mechanism for corrosion of iron-containing metals is electrobiocorrosion, the direct electron transfer from Fe^0^ to microbes to support anaerobic respiration (4). The concept of electrobiocorrosion was first proposed in studies of isolates from an enrichment culture in which pure Fe^0^ was provided as the sole electron donor (14). Two sulfate-reducing bacteria, now known as *Desulfovibrio ferrophilus* and *Desulfopila corrodens*, and the methanogen designated *Methanobacterium* strain IM1, corroded Fe^0^ faster than isolates not recovered from corrosion enrichments. However, there is no evidence that the direct electron transfer of electrobiocorrosion would necessarily be faster than corrosion in which H_2_ is an intermediary electron carrier. *D. ferrophilus*, *Ds. corrodens*, and strain IM1 are all capable of using H_2_ as an electron donor to support anaerobic respiration. When H_2_ production is the corrosion mechanism, factors such as a strain’s capability for attachment to metal surfaces, H_2_ affinities, and the potential for the release of hydrogenases or metabolites that might accelerate H_2_ production can influence corrosion rates (3, 4, 13). For example, studies with H_2_-utilizing methanogens demonstrated that the isolates that most rapidly corroded Fe^0^ had genes for extracellular hydrogenases to promote H_2_ formation from Fe^0^ (11). A rigorous approach to evaluate whether H_2_ is an intermediary electron carrier between Fe^0^ and microbes associated with corrosion is to delete hydrogenase genes to prevent the possibility of H_2_ serving as an electron donor (9, 15). Such gene deletion studies were not done with *D. ferrophilus*, *Ds. corrodens*, or strain IM1.

Electrobiocorrosion has been rigorously demonstrated with *Geobacter*, *Shewanella*, and *Methanosarcina* species. In studies with strains naturally unable to use H_2_ (*G. metallireducens*, *M. acetivorans*) or strains in which hydrogenases were deleted to prevent H_2_ uptake (*G. sulfurreducens*, *S. oneidensis*), deletion of genes for outer-surface, multi-heme *c*-type cytochromes inhibited corrosion (16–20). In each strain the cytochromes required for effective corrosion were previously demonstrated to participate in other forms of extracellular electron exchange.

When tools for genetic manipulation are not available, a comparison of corrosion with pure Fe^0^ versus 316L stainless steel (hereafter referred to simply as stainless steel) can provide insight into whether microbes are capable of electrobiocorrosion. Pure Fe^0^ abiotically oxidizes with the production of H_2_ at circumneutral pH (16, 21), whereas stainless steel does not produce H_2_ (17).

*Geobacter* species, which are capable of electrobiocorrosion, readily utilize stainless steel as an electron donor (17). However, microbes that rely on a H_2_ intermediate cannot. For example, *Desulfovibrio vulgaris* could reduce sulfate with pure Fe^0^ as the sole electron donor, but a hydrogenase mutant incapable of H_2_ uptake could not (15). Reliance on a H_2_ intermediate was also apparent from the parental, H_2_-utilizing strain’s inability to reduce sulfate with stainless steel as the electron donor (15).

Similar to *D. vulgaris*, *D. ferrophilus* and *Ds. corrodens* reduced sulfate with pure Fe^0^ as the electron donor, but not with stainless steel (22). These results indicated that *D. ferrophilus* and *Ds. corrodens* rely on H_2_ as an intermediary electron carrier from Fe^0^ to cells and are not capable of electrobiocorrosion. Similar studies with the methanogen *Methanobacterium* strain IM1 were not previously reported.

Studies on the corrosion mechanisms of a diversity of methanogens and acetogens are warranted because these physiological groups are often associated with corroding metal and several pure cultures accelerate corrosion (4, 11, 14, 23–30). Electrobiocorrosion is most expected in methanogens and acetogens that have the capacity for electron exchange with other extracellular donors or acceptors. For example, electrobiocorrosion has been demonstrated for *M. acetivorans* (19), a Type II *Methanosarcina* species (31) known for its ability to reduce extracellular electron acceptors (32, 33) and accept electrons for carbon dioxide reduction via direct interspecies electron transfer (DIET) (34). Unlike Type II *Methanosarcina*, Type I *Methanosarcina* can use H_2_ as an electron donor, and lack the outer-surface multi- heme cytochrome MmcA that is required for extracellular electron exchange in *M. acetivorans* (31). *M. barkeri,* a Type I *Methanosarcina* species, is capable of receiving electrons via DIET (35) and possibly electrodes (36), suggesting that it might also be capable of electrobiocorrosion. However, Fe^0^ oxidation under conditions that rule out the possibility of H_2_ serving as an intermediary electron carrier are required to more fully evaluate this possibility. *Methanothrix harundinaceae* (37) *Methanothrix thermoacetophila* (38) and *Methanothrix soehngenii* (39) are also capable of accepting electrons via DIET despite a lack of outer-surface cytochromes. Their potential for electrobiocorrosion has also not been previously reported.

It is also not clear whether acetogens that lack obvious outer-surface electrical contacts are able to directly accept electrons from extracellular electron donors (40). Negatively poised cathodes support the reduction of carbon dioxide to acetate by acetogens such as *Sporomusa ovata* and *Clostridium ljungdahlii* at potentials too positive to support abiotic H_2_ production, which was interpreted as direct electrode-to-microbe electron transfer (41, 42). However, subsequent studies have proposed that H_2_ is an intermediary electron carrier (9, 43). Studies with strains unable to use H_2_ have yet to be conducted to resolve this question.

Acetogenic *Clostridium* and *Sporomusa* and methanogenic *Methanothrix*, *Methanosarcina*, and *Methanobacterium* species frequently co-exist in corrosive biofilms (23, 26–28, 30, 44–46). It has been proposed that acetogens generating acetate from Fe^0^ oxidation supply acetate as an energy substrate for acetotrophic methanogens and as a carbon source for hydrogenotrophic methanogens (30, 46). Alternatively, acetogens and methanogens may compete for Fe^0^ (26, 44).

Here we report on the electrobiocorrosion potential of a diversity of methanogens and acetogens evaluated by comparing their ability to corrode pure Fe^0^ and stainless steel. The results expand the known phylogenetic and physiological diversity of electrobiocorrosive microbes and demonstrate that the methanogen strain IM1 is unlikely to participate in electrobiocorrosion.

## Materials and Methods

### Cultures and routine maintenance

All cultures were grown under strict anaerobic conditions in an N_2_:CO_2_ atmosphere (80:20, vol/vol). *Methanosarcina vacuolata* DH-1 was routinely cultured at 37 °C in defined mineral MA medium as previously described (31, 34) with methanol (20 mM) and acetate (40 mM) provided as substrates for growth. *Geobacter metallireducens* GS-15 (ATCC 53774) was routinely cultured anaerobically at 30°C in the previously described freshwater medium (47) with ethanol (20 mM) as the electron donor and Fe(III) citrate (56 mM) as the electron acceptor. *Sporomusa ovata* (DSMZ 2662) was grown with H_2_ (100 kPa) as the electron donor at 30°C in the DSMZ-recommended growth medium (DSMZ 311) with yeast extract, betaine, casitone, and resazurin omitted (42) and cysteine:sulfide (1 mM:0.5 mM) added from a 100-fold concentrated stock solution as a reductant. *Clostridium ljungdahlii* (DSM 13528) was grown anaerobically at 37 °C in DSMZ 879 medium with yeast extract and resazurin omitted. The 879 medium was also supplemented with cysteine:sulfide (1 mM:0.5 mM) and 10 mM sodium bicarbonate added from 100X anoxic stock solutions and H_2_ (100 kPa) was provided as the electron donor.

*Methanothrix soehringii* E1 was grown on DSMZ 334 medium at 37 °C with 60 mM acetate provided as a substrate for growth as previously described (39).

*Methanobacterium* strain IM1 was cultured on DSMZ 334 modified with the following additions: 18.3 g/L NaCl, 0.33 g/L CaCl_2_, 6.1 g/L MgCl_2_, 0.09 g/L KBr, 0.006 mg/L Na_2_SeO_3_ 5 H_2_O, 0.008 mg/L Na_2_WO_4_ 2 H_2_O, 13.4 mM NaHCO_3_, and 1 mM cysteine-HCl, and 0.5 mM NaS 9 H_2_O. Incubation was at 30 °C with H_2_ as the electron donor (100 kPa).

### Growth with pure Fe^0^ or stainless steel as potential electron donors

Pure Fe^0^ granules (10 g; 1-2 mm; Thermo Scientific) or 316 L stainless steel (12 pieces; 2 mm × 3 mm × 3 mm; Institute of Metal Research, Chinese Academy of Sciences) were pretreated as previously described (16, 17) and added to 50 ml of medium in 156 ml serum bottles under N_2_:CO_2_ (80:20, vol/vol). When noted, 1 mM sodium acetate was added for the studies with *M. vacuolata* and *Mx. soehngenii*.

To evaluate the possibility that cells might release hydrogenases or other factors that promote H_2_ production from Fe^0^, supernatants of cultures grown on pure Fe^0^ were anaerobically filtered through sterile 0.2 µm syringe filters (Corning Inc, NY). The culture filtrates (10 ml) were added to sterile tubes containing 2 g of pure Fe^0^ granules and H_2_ production was compared with granules and sterile medium.

### Evaluation of strain IM1 DIET

Before DIET between strain IM1 and *Geobacter metallireducens* could be tested, pure cultures of both strains needed to adapt to grow at similar salt concentrations. Both strains were separately cultured on modified 334 IM1 medium described above with 4 g/L NaCl, 0.15 g/L CaCl_2_, and 1.3 g/L Mg/Cl_2_. Ethanol (20 mM) and Fe(III) citrate (56 mM) were provided as electron donor and acceptor for pure cultures of *G. metallireducens* and H_2_ (100 kPa) and CO_2_ were provided as the electron donor and acceptor for strain IM1 during adaptation experiments. After adaptation, 0.5 ml of both adapted strains was inoculated into 50 ml of IM1 modified 334 medium with ethanol (20 mM) provided as the electron donor and CO_2_ provided as the electron acceptor. Co-cultures were then grown at 30°C.

When noted, granular activated carbon (GAC, Sigma, C2889, 8-20 mesh, various concentrations) or magnetite nanoparticles (10 mM) were added to the co-culture medium before autoclaving. The surface area and resistivity of the GAC used was 600-800 m^2^/g (dry basis) and 1375 μΩ-cm, respectively. Magnetite nanoparticles with diameters of 20-50 nm were prepared as previously described (48).

### Analytical techniques

Methane and H_2_ in headspace gas was measured with gas chromatography. H_2_was monitored with a thermal conductivity detector in a gas chromatograph (Agilent Technologies G1530A, USA) equipped with a CARBONXENTM 1010 PLOT column (30 m × 0.53 mm). The oven temperature was 40°C, and the detector temperature was set at 225°C. The carrier gas was N_2_. Methane was measured with a gas chromatograph (Shimadzu, GC-8A) with a flame ionization detector as previously described (49).

Dissolved acetate was measured with a SHIMADZU high performance liquid chromatograph (HPLC) with an AminexTM HPX-87H Ion Exclusion column (300 mm × 7.8 mm) and an eluent of 8.0 mM sulfuric acid. Dissolved ethanol was measured with a gas chromatograph (Clarus 600; PerkinElmer Inc., CA) equipped with a headspace sampler and a flame ionization detector, as previously described (50).

## Results and Discussion

### Electrobiocorrosion by Methanosarcina vacuolata, a Type I Methanosarcina

*Methanosarcina vacuolata* DH-1 is a Type I *Methanosarcina* species that in addition to utilizing H_2_, acetate, methanol, and methylamines for methane production, can accept electrons via DIET for the reduction of carbon dioxide to methane (31). *M. vacuolata* produced methane with pure Fe^0^ as the sole electron donor (Figure 1A).

**Figure 1.**
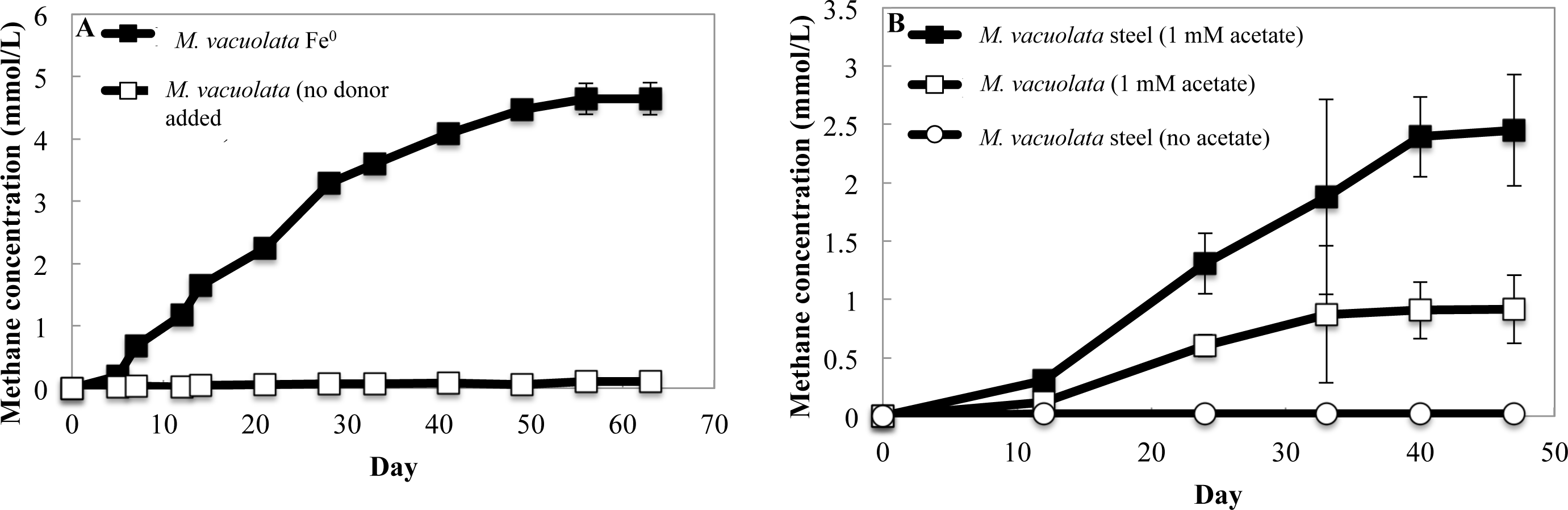
*M. vacuolata* DH-1 methane production with pure Fe^0^ or 316L stainless steel as the potential electron donor. (A) Pure Fe^0^ as the sole electron donor. (B) Stainless steel with or without 1 mM acetate added. The results are the means and standard deviation of triplicate cultures for each treatment. Significant differences in methane production were observed when cells were grown on pure Fe^0^ compared to a no donor control (p-value 6.03x10^-5^) and when cells were grown on stainless steel in the presence of 1 mM acetate compared to 1 mM acetate without stainless steel (p- value 0.0013).

Filtrates of *M. vacuolata* growing on Fe^0^ did not stimulate H_2_ production from Fe^0^ (Supplementary Figure S1A), indicating that *M. vacuolata* did not release hydrogenase or other factors promoting Fe^0^ oxidation with the reduction of protons.

*M. vacuolata* also produced methane with stainless steel as the electron donor, but only when 1 mM acetate was included in the medium (Figure 1B). Methane production substantially exceeded the maximum of 1 mmol/L methane that could have been generated from the added acetate, indicating that electrons for methane production were also derived from the stainless steel. *M. vacuolata* methane production with electrons derived from stainless steel are indicative of electrobiocorrosion because stainless steel does not produce H_2_ (17).

Previous studies demonstrated that *M. acetivorans* is also only capable of electrobiocorrosion when acetate is available (19). Electrobiocorrosion may not provide enough energy to support growth from carbon dioxide reduction to methane when it is the only source of electrons, or acetate may be required to balance metabolic fluxes either for biosynthetic or methane-generating pathways (19).

Production of methane by *M. vacuolata* from pure Fe^0^ in the absence of added acetate suggests that H_2_ was an intermediary electron carrier under those conditions. The ability of *M. vacuolata* to accept electrons from Fe^0^ either with H_2_ as an intermediary electron carrier, or via electrobiocorrosion, is similar to *S. oneidensis*, which is also capable of both processes (18).

### Methanothrix soehngenii electrobiocorrosion

Like other *Methanothrix* species, *Methanothrix soehngenii* strain E1, an anaerobic digester isolate, cannot use H_2_ as an electron donor, but converts acetate to methane and accepts electrons via DIET for carbon dioxide reduction to methane (39). *M. soehngenii* produced negligible methane when pure Fe^0^ was the sole electron donor (Figure 2A). However, when 1mM acetate was added as an additional energy source, methane was produced in excess of that possible from acetate alone (Figure 2A). These results are similar to previous studies on *M. acetivorans* electrobiocorrosion in which the H_2_ available from pure Fe^0^ could not be utilized, but electrobiocorrosion proceeded in the presence of acetate (19). *Mx. soehngenii* methane production from stainless steel followed a pattern similar to that with pure Fe^0^ (Figure 2B). Acetate was required, but the total amount of methane produced was much greater than could be attributed to acetate metabolism alone.

**Figure 2.**
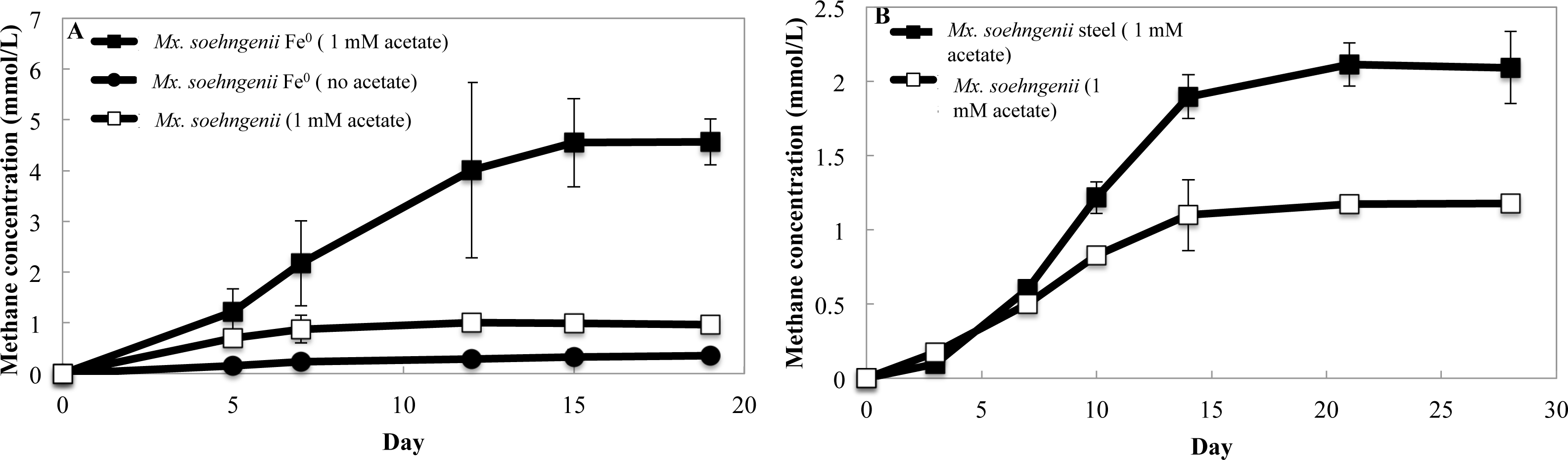
*Mx. soehngenii* E1 methane production with pure Fe^0^ or 316L stainless steel as the electron donor. (A) Pure Fe^0^ as the sole electron donor. (B) Stainless steel with or without 1 mM acetate added. The results are the means and standard deviation of triplicate cultures for each treatment. Significant differences in methane production were observed when cells were grown on Fe^0^ (p-value 0.002) or stainless steel (p-value 0.001) with 1 mM acetate compared to a growth on 1 mM acetate alone.

Thus, the current evidence suggests that Type I *Methanosarcina* and *Methanothrix* species, both of which lack outer-surface cytochromes, are capable of electrobiocorrosion. This is consistent with DIET studies in which Type I *Methanosarcina* and *Methanothrix* species directly accepted electrons from *Geobacter metallireducens* without a H_2_ intermediary (31, 35, 37–39, 51–53). The electrical contacts on the outer surface of Type I *Methanosarcina* and *Methanothrix* remain to be determined. Potential candidates include putative outer-surface proteins whose genes were highly expressed in *M. barkeri* and *Mx. thermoacetophila* during growth via DIET, including surface-associated quinoproteins (38, 51, 54).

### *Methanobacterium* strain IM1 lack of electrobiocorrosion

*Methanobacterium* strain IM1 was previously proposed to be capable of electrobiocorosion (14), but as detailed in the Introduction, the evidence supporting this conclusion is circumstantial and inconclusive (3, 4). Strain IM1 effectively produced methane from pure Fe^0^ (Figure 3A), generating 1.8 times (p-value=0.024) more methane than *M. vacuolata* from the same amount of Fe^0^ (Figure 1A). However, strain IM1 did not produce methane from stainless steel (Figure 3A).

**Figure 3.**
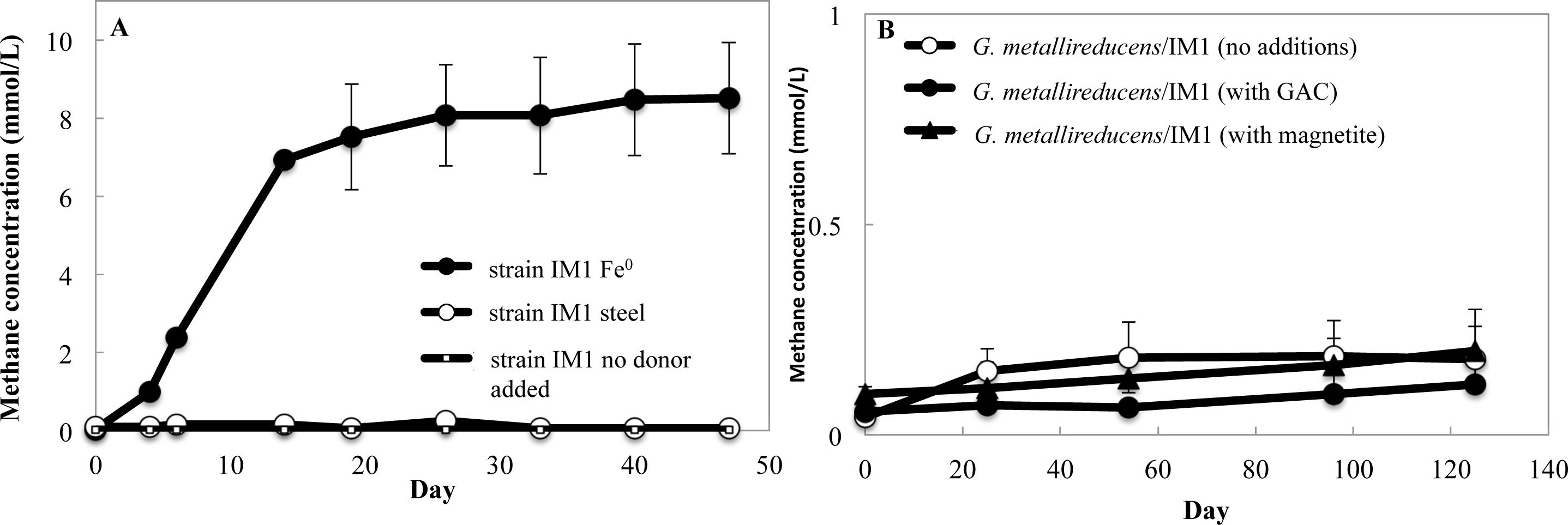
Evaluation of *Methanobacterium* strain IM1 electron uptake capabilities. (A) Methane production with pure Fe^0^ or stainless steel as the electron donor as well as in a control with no electron donor. (B) Lack of methane production in co-cultures of *G. metallireducens* and strain IM1 in the absence of conductive materials, in the presence of granular activated carbon (GAC; 2 g/50 ml), or in the presence of 10 mM magnetite. The results are the means and standard deviation of triplicate cultures for each treatment. Significant differences in methane production were observed when cells were grown on pure Fe^0^ (p-value 0.0008) compared to the no donor control.

Culture filtrates of strain IM1 did not produce H_2_ from Fe^0^ beyond that generated in abiotic controls (Supplementary Figure S1B), indicating that strain IM1 does not release extracellular hydrogenases. Development of tools for genetic manipulation of strain IM1 could further elucidate its corrosion mechanisms, but with the evidence currently available it appears that strain IM1 is not capable of electrobiocorrosion and relies on a H_2_ intermediary electron carrier to accept electrons from Fe^0^. The faster methane production of strain IM-1 (Figure 3A) than *M. vacuolata* (Figure 1A) during growth on pure Fe^0^ illustrates the fallacy of the assumption (14) that faster electron uptake is indicative of electrobiocorrosion. The distinct carbon and hydrogen stable isotope fractionation displayed by strain IM1 during growth with H_2_ and Fe^0^ (55) may, therefore, reflect different kinetic constraints of H_2_ uptake, which translates as mass transfer limitations, known to affect the extent of isotope fractionation (56).

To further examine strain IM-1’s electron uptake capabilities, attempts were made to determine whether it was able to accept electrons donated from direct interspecies electron transfer (DIET). Co-culture growth with *G. metallireducens* metabolizing ethanol as the electron donating partner is possible via DIET, but not with a H_2_ intermediary because *G. metallireducens* does not produce H_2_ under these conditions (57–59). *G. metallireducens* and strain IM1 were adapted to grow well separately in the same medium, but with ethanol (*G. metallireducens*) or H_2_ (strain IM1) as the electron donor and Fe(III) citrate (*G. metallireducens*) or CO_2_ (strain IM1) as electron acceptor. Co-cultures with ethanol as the electron donor were initiated with adapted *G. metallireducens* and IM1 strains, but no growth was observed after 120 days, even in cultures supplemented with conductive materials such as granular activated carbon (60) or magnetite (48) that are known to promote DIET (Figure 3B). These results further support the conclusion that strain IM1 is not capable of direct electron uptake from extracellular electron donors.

### Electrobiocorrosion with *S. ovata* but not *C. ljundahlii*

Acetogens such as *Sporomusa* and *Clostridium* species are often associated with corroding iron metals (9, 26, 30, 44). *S. ovata* produced substantial acetate with pure Fe^0^ as the sole electron donor (Figure 4A). This result contrasts with the previous finding that *S. ovata* is unable to effectively utilize Fe^0^ as an electron donor for acetogenesis (13, 30). Less acetate was generated with stainless steel as the source of Fe^0^, but acetate levels were consistently higher than in controls without an added electron donor (Figure 4A). The production of acetate from stainless steel, which is not known to release H_2_ (17), suggests that *S. ovata* is capable of directly accepting electrons from Fe^0^. *S. ovata* culture supernatant incubated with pure Fe^0^ did not produce more H_2_ than abiotic controls, indicating that *S. ovata* does not secrete extracellular hydrogenases (Supplementary Figure S1C). However, development of genetic tools to further evaluate this possibility is warranted to help better understand the potential role of H_2_ during *S. ovata* corrosion (13, 29, 43, 61) and to identify possible direct electron uptake mechanisms.

**Figure 4.**
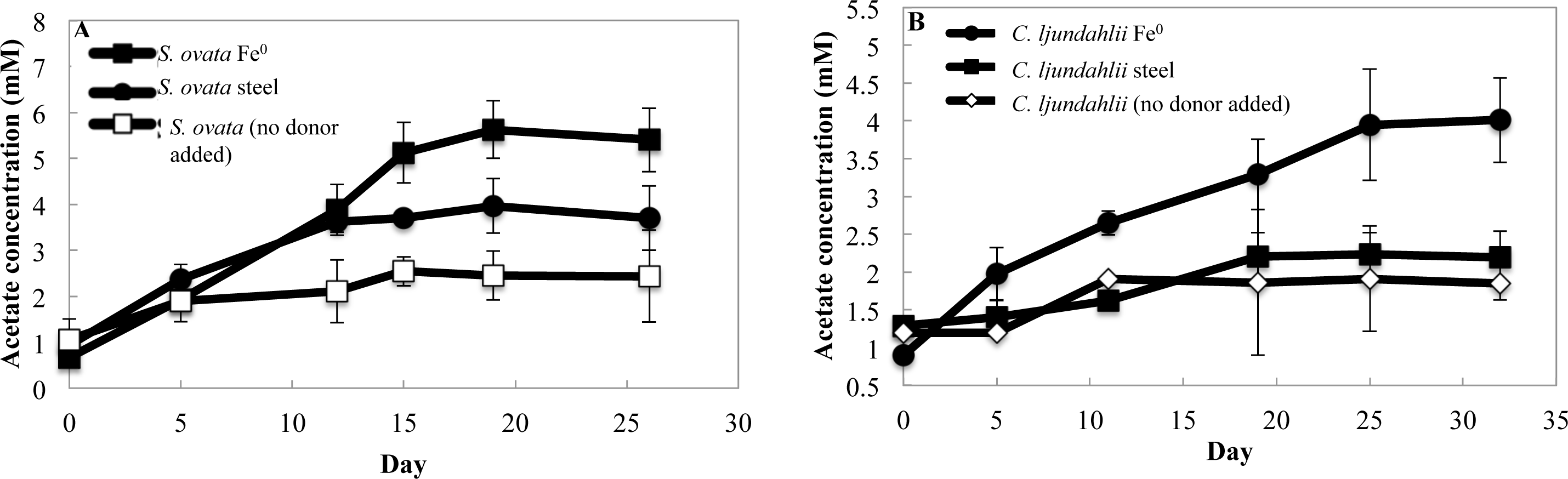
Acetate production in cultures of *S. ovata* or *C. ljundahlii* with pure Fe^0^ or stainless steel as the potential electron donor. (A) *S. ovata* acetate production. Significant differences were observed when cells were grown on pure Fe^0^ (p-value 0.0003) or stainless steel (p-value 0.0005) compared to no donor controls. (B) *C. ljundahlii* acetate production. Significant differences were observed when cells were grown on pure Fe^0^ (p-value 0.0006), but not stainless steel (p-value > 0.05) compared to no donor controls. The results are the means and standard deviation of triplicate cultures for each treatment.

As with *S. ovata*, the question of whether or not *Clostridium ljundahlii* can directly accept electrons from electrodes is unresolved (13, 62). Microbial electrosynthesis with *Clostridium ljungdahlii* benefits from H_2_ mediating electron transfer between electrodes and cells (40, 62). *C. ljundahlii* readily produced acetate with pure Fe^0^ as the electron donor (Figure 4B). However, acetate concentrations were not significantly higher than the no-electron donor control when stainless steel was the electron donor (Figure 4B). These results indicate that the role of *C. ljundahlii* in iron corrosion is probably restricted to H_2_ consumption. Filtrates of *C. ljundahlii* did not produce significantly more H_2_ than abiotic controls (Supplementary Figure S1D).

### Acetogen acetate production enables methanogenesis

It has previously been proposed that acetate produced by acetogens from Fe^0^ corrosion can support the growth of acetotrophic methanogens associated with corrosion (27, 28). Alternatively acetogens and methanogens may compete for Fe^0^ as an electron donor (26, 44). The findings reported here and previously (19) that *Methanosarcina* and *Methanothrix* species are capable of electrobiocorrosion in the presence of acetate suggested that acetogens might facilitate methanogen electrobiocorrosion. This possibility was evaluated with co-cultures of *S. ovata* and *M. soehngenii.* With H_2_ as the electron donor, *S. ovata* produced acetate, and *Mx. soehngenii*, which is incapable of H_2_ consumption, consumed the acetate with the production of methane (Figure 5A and 5B). The presence of *Mx. soehngenii* appeared to enhance *S. ovata* acetate production because even though acetate was a transitory intermediate in the co-culture, the maximum acetate that accumulated (19.17 ± 1.62 mM) was significantly higher than the maximum acetate generated (12.66 ± 0.52 mM) when *S. ovata* alone was grown on H_2_ (Supplementary Figure S2).

**Figure 5.**
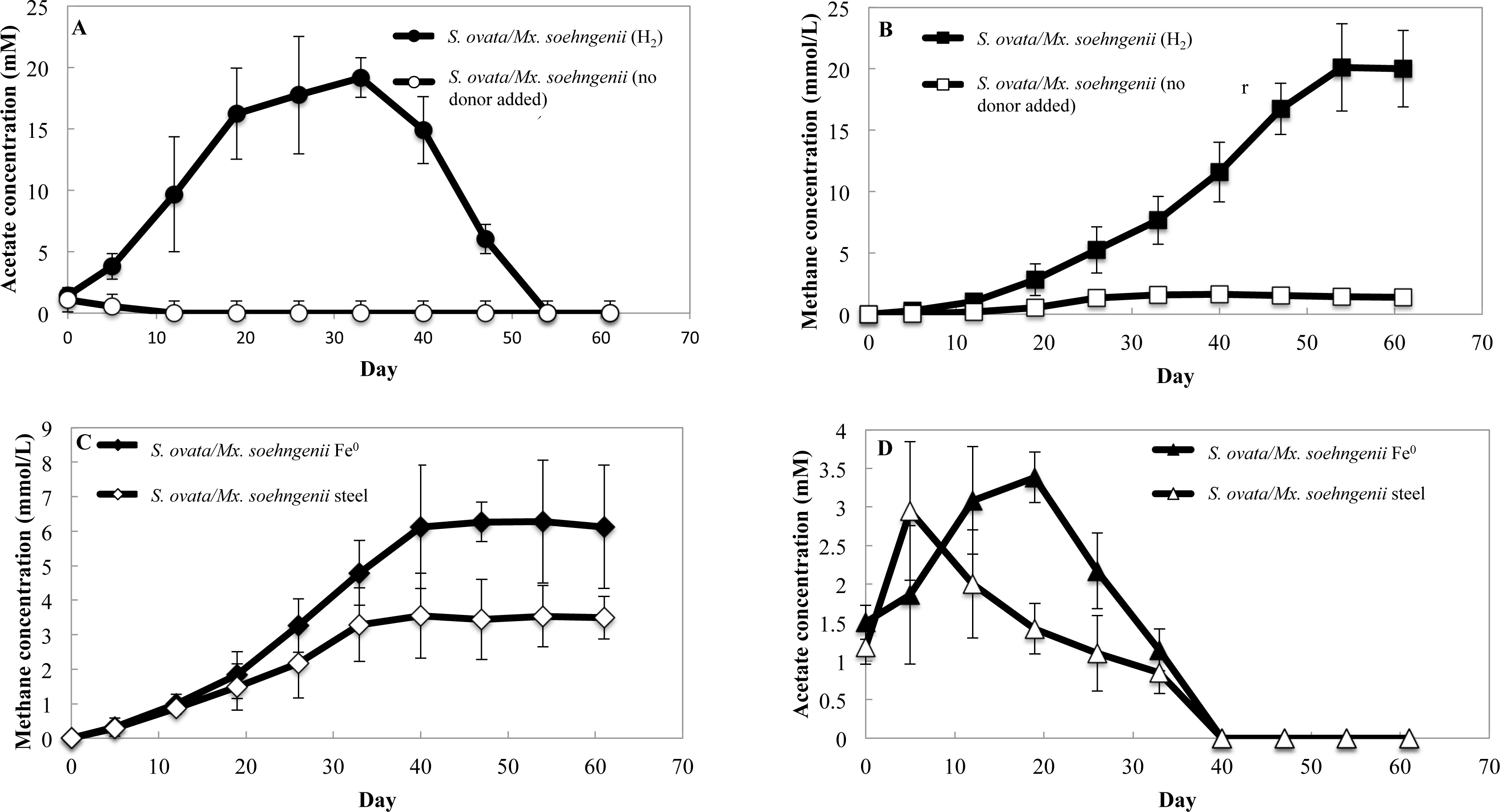
*S. ovata/Mx. soehngenii* co-culture growth with H_2_, pure Fe^0^, or steel as the electron donor. Acetate concentrations (A) and methane production (B) of the co-culture growing with H_2_ as the electron donor. Methane production (C) and acetate concentrations (D) of the co-culture growing with pure Fe^0^ or steel as the electron donor. The results are the means and standard deviation of triplicate cultures for each treatment.

Although *Mx. soehngenii* was unable to produce methane with pure Fe^0^ or stainless steel as the electron donor in the absence of added acetate (Figure 2), methane was produced when *Mx. soehngenii* was cocultured with *S. ovata* (Figure 5C). Acetate only transiently accumulated (Figure 5D) as *Mx. soehngenii* converted it to methane. The amount of methane the co-culture produced from pure Fe^0^ (6.28 ± 0.57 mM) or stainless steel (3.55 ± 0.89 mM) was not significantly different from the amount of acetate that *S. ovata* alone (Figure 4A) produced from pure Fe^0^ (5.62 ± 0.70) or stainless steel (3.97 ± 0.66). Thus, with the data available it is not possible to determine whether *Mx. soehngenii* was solely producing methane from the acetate that *S. ovata* generated or was also directly contributing to Fe^0^ oxidation via electrobiocorrosion.

### Implications

The results demonstrate that several anaerobes that lack the outer-surface *c*- type cytochromes shown to be important for electrobiocorrosion in other microbes (16–20) are also capable of direct electron uptake from Fe^0^. Electrobiocorosion was most apparent with *Mx. soehngenii*, in which a role for H_2_ could be eliminated because *Mx. soehngenii* is unable to utilize H_2_ (39, 63, 64). The studies with stainless steel, which does not generate H_2_ (17), further confirmed the bioelectrocorrosion capabilities of *Mx. soehngenii* and indicated that *M. vacuolata* and *S. ovata* could also accept electrons from Fe^0^ without a H_2_ intermediate. Although *Methanobacterium* strain IM1 was among the initial cohort of microbes first proposed to be capable of electrobiocorrosion (14) strain IM1 was unable to use stainless steel as an electron donor, indicating that it relies on H_2_ produced from Fe^0^ as its electron donor. These results are similar to those previously reported for *D. ferrophilus* and *Ds. Corrodens* the other cohort members (22) and further emphasize that corrosion mechanisms cannot be discerned based simply on relative rates of corrosion (3, 13).

The expanding known diversity of microbes capable of electrobiocorrosion suggests that this process should be considered as a possibility in most instances of microbial corrosion. Needed are studies of natural communities actively corroding iron-containing surfaces at a sufficient level of detail to discern the relative roles of electrobiocorrosion, H_2_ uptake, and more indirect process such as the production of sulfide or acids, in corrosion (3, 4).

