## Supplementary Figures for "Electrobiocorrosion by Microbes without Outer-Surface Cytochromes"

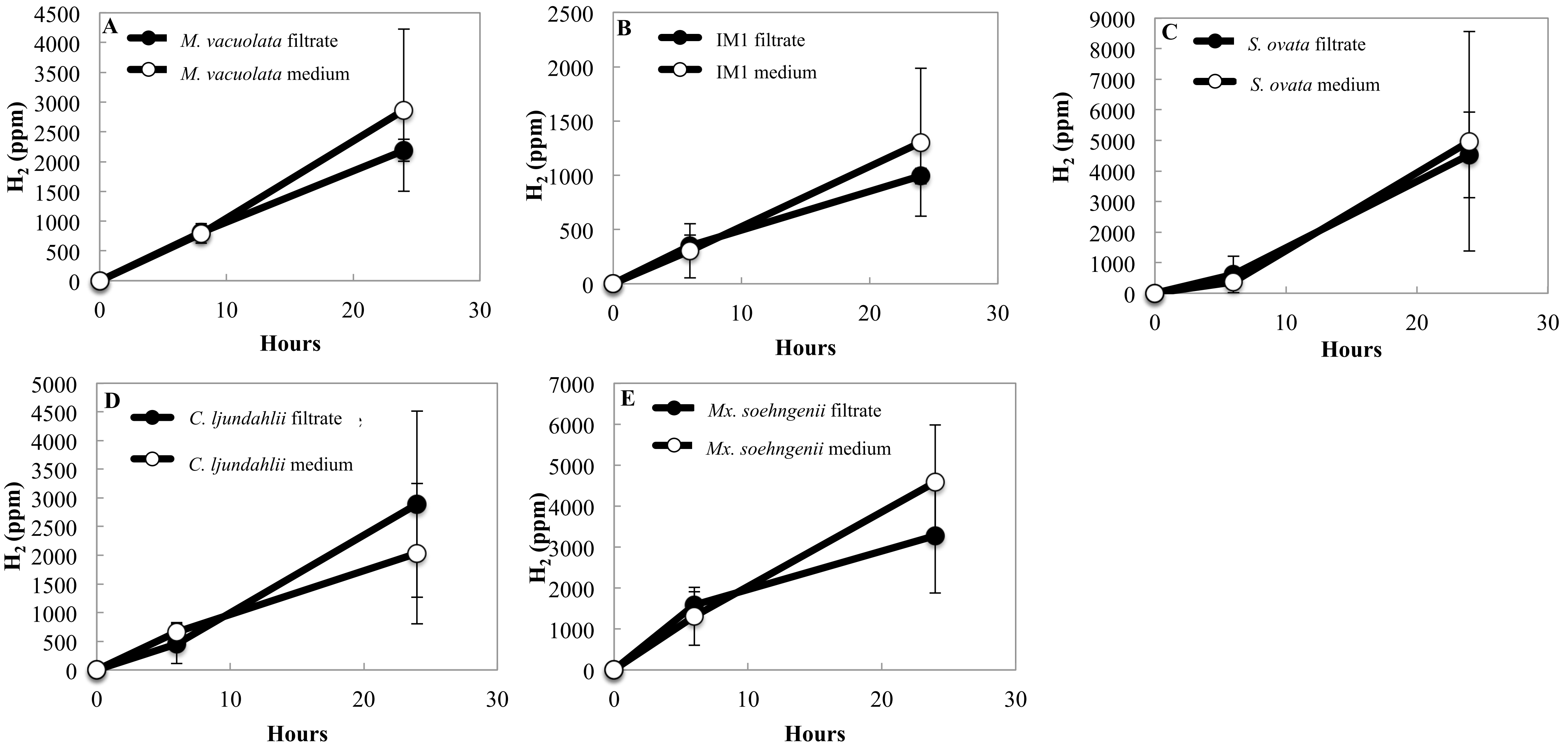


Supplementary Figure S1. Cathodic $H_2$ production by pure $Fe^0$ in the presence or absence of filtered supernatant from the various cultures. For filtrate samples: one milliliter of the appropriate mid-logarithmic culture was filter sterilized and inoculated into 9 ml of medium with 2 g pure $Fe^0$ granules. For control samples: one ml of un-inoculated medium was added to 9 ml of medium with 2 g pure $Fe^0$ granules. Error bars represent triplicate samples.

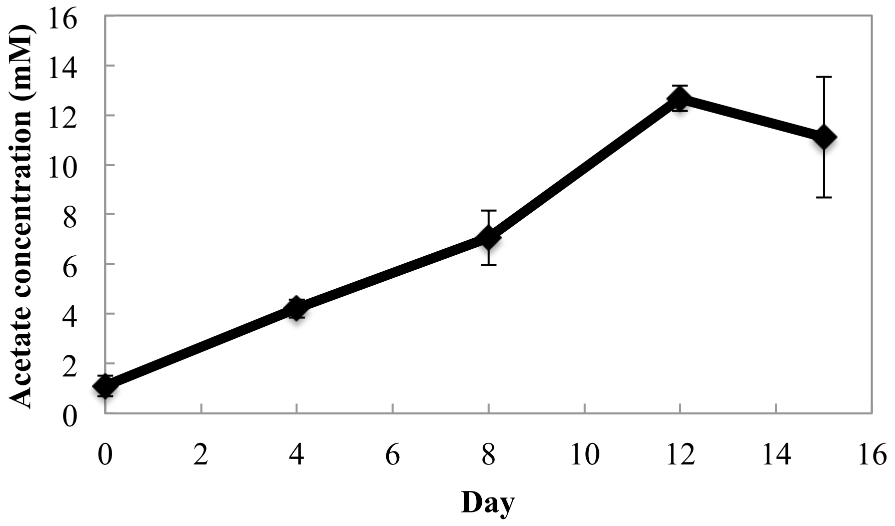


Supplementary Figure 2. Acetate generated by *Sporomusa ovata* cultures grown with $H_2$ (100 kPa) as the sole electron donor and $CO_2$ as the sole electron acceptor. The results are the means and standard deviation of triplicate cultures.
